# Common Vetch: a drought tolerant, high protein neglected leguminous crop with potential as a sustainable food source

**DOI:** 10.1101/2020.02.11.943324

**Authors:** Vy Nguyen, Samuel Riley, Stuart Nagel, Ian Fisk, Iain R. Searle

## Abstract

Global demand for protein is predicted to increase by 50% by 2050. To meet the increasing demand whilst ensuring sustainability, protein sources that generate low-greenhouse gas emissions are required and protein-rich legume seeds have the potential to make a significant contribution. Legumes like common vetch (*Vicia sativa*) that grow in marginal cropping zones and are drought tolerant and resilient to changeable annual weather patterns, will be in high demand as the climate changes. In common vetch, the inability to eliminate the γ-glutamyl-β-cyano-alanine (GBCA) toxin present in the seed has hindered its utility as a human and animal food for many decades, leaving this highly resilient species an “orphan” legume. However, the availability of the vetch genome and transcriptome data together with the application of CRISPR-Cas genome editing technologies lay the foundations to eliminate the GBCA toxin constraint. In the near future, we anticipate that a zero-toxin vetch variety will become a significant contributor to global protein demand.

## Introduction

Global demand for protein is predicted to increase by a staggering 50% by 2050 (Westhoek et al., 2011; Henchion et al., 2017). With an increasing global population and increasing demand for animal-derived protein, the sustainability of agriculture systems has been brought into question. Over the last two centuries, the expanding livestock industry has led to significant deforestation and overgrazing of natural grassland environments such that it has caused decreased terrestrial biodiversity and increased greenhouse gas emissions and contributed to climate change and global warming (Henchion et al., 2017). In order to meet the increasing protein demand and protect our environment, more sustainable protein food sources are required. Cheap plant-based protein, such as legume seeds, represent an environmentally sustainable option that is well suited for developing countries with rapidly growing populations (Asgar et al., 2010). Moreover, to cope with increasingly unpredictable climate change and expansion of marginal cropping areas, breeding strategies for more drought tolerant and resilient crops will be vital (Sivakumar et al., 2005; Lobell and Gourdji, 2012). A legume that could be exploited for this scenario is the common vetch (*V. sativa*). Common vetch is able to grow in marginal cropping zones whilst being resilient to variable annual weather patterns mainly through superior drought tolerance (White et al., 2005). One study demonstrated that vetch could withstand water deficit for up to 24 days and show full restoration of biotic function once regular watering had resumed (Tenopala et al., 2012). Drought and heat tolerant crops are increasingly desirable in the face of rising global temperatures and increasingly prolonged periods of drought brought on by climate change (Mba et al., 2018).

For many decades, the inability to remove the γ-glutamyl-β-cyano-alanine (GBCA) seed toxin has hindered common vetch’s use in agriculture (Pfeffer and Ressler, 1967; Roy et al., 1996), leaving this resilient plant as an “orphan” legume. We envisage that the development of a zero-toxin vetch variety would facilitate its use for animal feed, specifically chickens and pigs, and human consumption (Ressler et al., 1997; Collins et al., 2002). As the production costs of common vetch are approximately 50% less than competing legumes such as lentils (unpublished data), we predict that zero-toxin varieties would rapidly surpass lentil in pig and poultry production. Zero-toxin common vetch will immediately generate new domestic markets, such as feed for the poultry industry, but it will also open new export markets for Australia and other countries thereby increasing export revenue and increasing farm profitability and indirectly increase investment for their local communities.

### Common Vetch: a versatile pasture crop that provides multiple benefits for the farm

Common vetch which is shown in Figure 1 belongs to the *Fabaceae* (legume) family, within the genus *Vicia*. This genus contains about 140 species including woolly-pod vetch (*V. villosa*) and broad bean (*V. faba*). Other *Fabaceae* genera also contain so-called vetches; of which two examples are *Astragalus* (containing the milkvetches) and *Lathyrus* (containing *L. ochrus*, the cyrpus-vetch). Nowadays, common vetch is commonly found both in natural and agricultural settings across Europe, Asia, North America, some parts of South America, Africa, the Mediterranean and Australia (Navrátilová et al., 2003; Ford et al., 2008).

**Figure 1.**
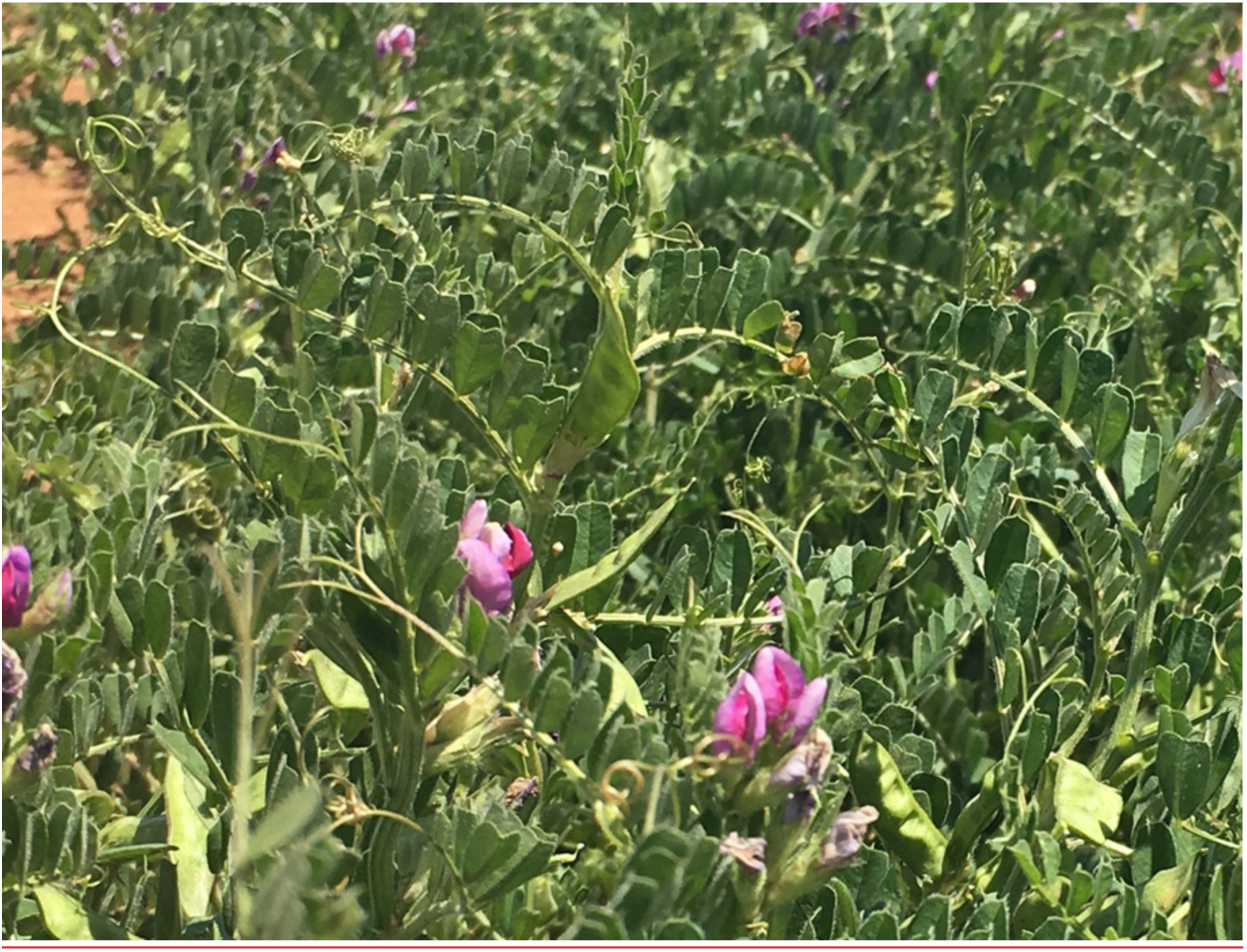
Field grown *Vicia sativa* growing at the National Vetch Breeding Program in South Australia (2017).

Like other legumes, common vetch forms a symbiosis with nitrogen-fixing bacteria (*Rhizobia*) that fix atmospheric nitrogen into nitrogenous compounds available to the plant, hence reducing the need for application of expensive nitrogen fertilizer and subsequent rotation crops. Often, common vetch is used as a green manure which, when incorporated into the soil, provides valuable carbon and nitrogen for rotation crops such as wheat and barley. Additional soil carbon often increases water-holding capacity and ability to bind nutrients including nitrate (Reeves, 1997; Bünemann et al., 2018). Furthermore, common vetch biomass can also be used for forage, fodder, pasture, silage or hay and the seed may safely be used as a protein-rich feed component for ruminant animals (Enneking, 1995). Common vetch is well suited as a pasture species as it forms many adventitious shoots that are either buried or close to the soil surface thus giving it the ability to be resilient to heavy grazing (Rathjen, 1997).

Despite common vetch’s versatile uses, the production of vetch is still limited. Data collected by the Food and Agriculture Organisation (FAO) in 2017 showed that, globally, the area harvested and the production of vetch were about 0.6 million ha and 0.9 million tonnes, respectively. Vetch occupied about 0.3% of land usage and accounted for only 0.2% of the production of major legume group. This is 12 times less than the area harvested and 8 times less than the production of lentils (Figure 2). Common vetch’s limited production is mainly attributed to the anti-nutritional compounds existing in the seeds which will be further discussed in the next section.

**Figure 2.**
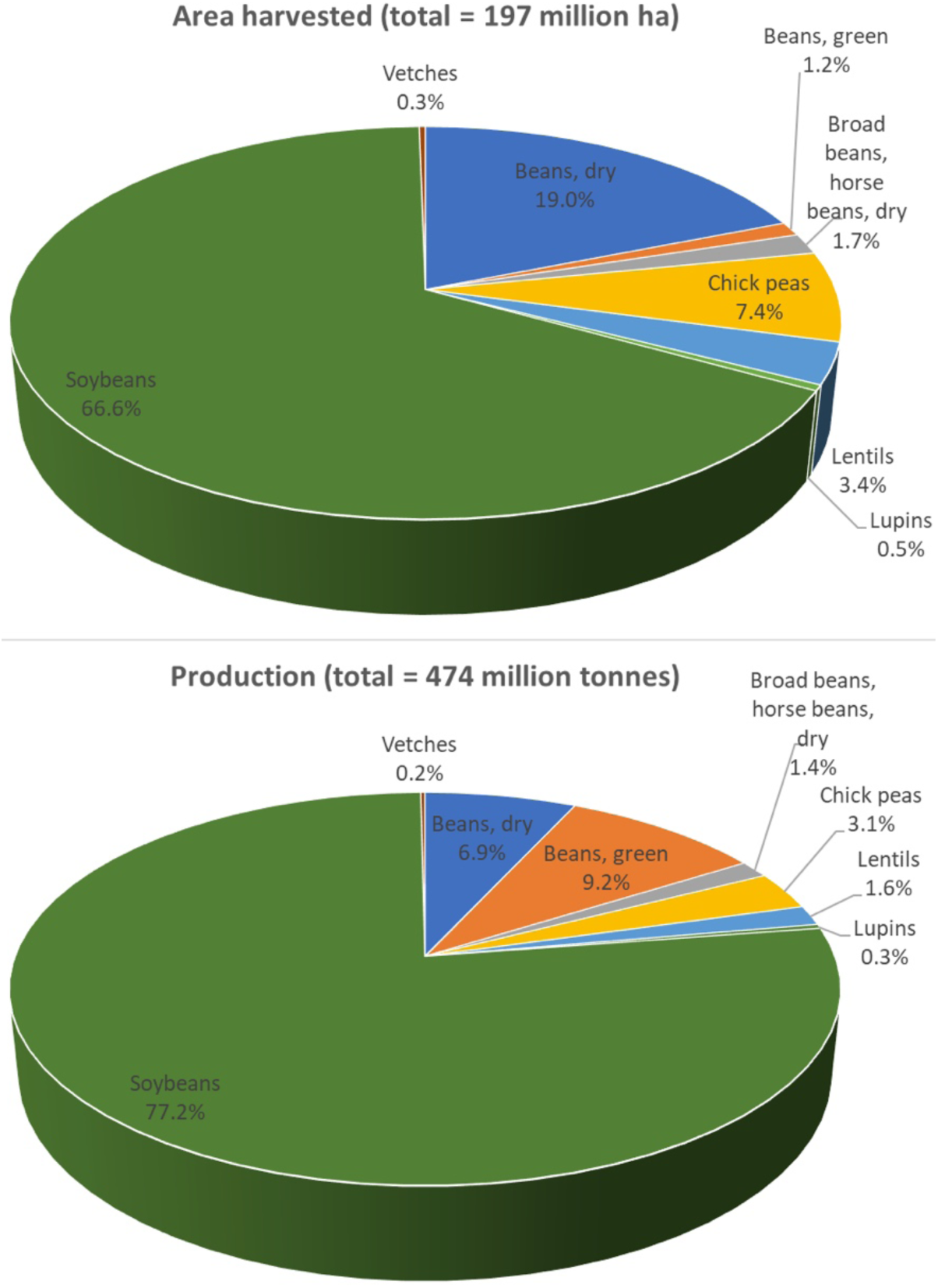
Common vetch production when compared to other legumes globally in 2017. In general, common vetch occupied about 0.3% of the area harvested and 0.2% of the production. For vetch, total area harvested in 2017 was 560,077 ha and total production was 920,536 tonnes. Data was published by the Food and Agriculture Organisation (FAO) (http://www.fao.org/).

### Common Vetch seeds: important nutritional attributes

Common vetch seed appeared in the diets of hunter-gatherers as early as 12,000 – 9,000 BP as evident in archaeobotanical analysis of samples from the Santa Maira cave in Alicante, Spain (Aura et al., 2005; Mikić, 2016). Today, common vetch is globally distributed and its spread is thought to have occurred through the inadvertent selection and trading of vetch seeds as a weedy contaminant with other legume seeds (Erskine et al., 1994). This has led to the suggestion that selection of common vetch amongst other palatable legumes, such as lentils, lead to vegetative and seed mimicry of vetch to lentils (Erskine et al., 1994). Due to the similarity in the seeds between vetch and lentil, there was a period of time when the vetch variety “blanche fleur” was inappropriately promoted as an inexpensive alternative to lentils. However, Tate and Enneking (1992) raised the issue of vetch toxicity and indicated that vetch seeds were unsuitable to be eaten by monogastric animals, including humans. Since then, vetch has been restricted to use in pastures as the seeds can be safely eaten by ruminants and some monogastric animals, such as chicken (less than 40%) and pig (less than 20%) in low amounts (Rathjen, 1997).

Compared with other legumes, common vetch exhibits high concentrations of crude protein between 24 to 32% (Francis et al., 2000) and carbohydrates and fiber are comparable to that of lupin (Valentine and Bartsch, 1996). Common vetch seeds contain eighteen amino acids and the ratio of essential amino acids/non-essential amino acids is about 0.7 (Mao et al., 2015) which is significantly higher than the 0.38 recommended by WHO (Who, 2007). The principle essential amino acids are arginine and leucine at average concentrations of 2.4% and 2.1%, respectively. Glutamic and aspartic acids are the predominant non-essential amino acids within common vetch, averaging levels of 5.5% and 3.7%, respectively (Mao et al., 2015). The species also contains a much lower proportion of lipids (only 1.5 – 2.7%) compared with soybean and this is primarily composed of unsaturated fatty acids (Mao et al., 2015). Presently, common vetches are mainly used as animal feed for ruminants and it can be argued that higher value returns could be observed through the repurposing of common vetch as a human food crop. Obstructing the development of the higher value product is a range of anti-nutritional factors, the most significant of which are the dipeptide γ-glutamyl-β-cyanoalanine (GBCA) and the free amino acid β-cyano-L-alanine (BCA), that exist in relatively high concentrations in the seed at approximately 2.6% and 0.9%, respectively (Tate et al., 1999). Such compounds are toxic to monogastric species, such as chickens or pigs, but have no obvious effect upon ruminant species, including beef cattle (Valentine and Bartsch, 1996). In addition, vicine, convicine, and other anti-nutritional factors typical of leguminous crops, including tannins, trypsin inhibitors and polyphenols are also present. To fully utilize the nutrition of common vetch, these also require reduction after the elimination of GBCA and BCA. Currently, the proportion of common vetch seed within feed that can be safely consumed without deleterious effects is 10% for piglets and 20% for adult pigs. For chickens it is higher at about 40% (Harper and Arscott, 1962; Rathjen, 1997).

### Limitation of GBCA detoxification using conventional methods

Without the toxic compounds in the seeds, vetch would be highly nutritious. Therefore, a number of methods have been investigated to remove the toxin. Post-harvest processing efforts to lower the GBCA toxin levels within the seed have previously involved simple soaking, continuous flow through soaking and boiling methods (Rathjen, 1997). The seed soaking method alone was insufficient to lower GBCA levels as consumption of soaked seed during feeding trials by chickens reduced egg production, daily food consumption and feed conversion ratios were also diminished (Farran et al., 1995). In contrast, boiling the seed produced lower toxin levels such that it could be included in chicken feed at levels of up to 25% without negative effects on growth rates (Kaya et al., 2013). However, consumption of boiled vetch seed that had the broth periodically discarded during boiling resulted in 20% reduced growth rates in chickens when compared with conventional feed with similar protein amounts (Ressler et al., 1997). This boiling method combined with periodically discarded water had 45% decreased seed mass as water soluble vitamins, like vitamin B, water soluble proteins and carbohydrates were leached during processing (Ressler et al., 1997) and this correlated with the reduced chick growth rate. Autoclaving the seed as a processing method has also been investigated and assessed in feeding trials of laying hens and dairy cows. In the dairy cow, it was found that the seed autoclaved at 120°C for 30 minutes resulted in removal of all the thermally sensitive toxins, whilst most of the nutritional content was retained (Aguilera et al., 1992). In the laying hen trial, overall growth rate was found not significantly different between animals fed with autoclaved or raw vetch seed suggesting that the non-heat labile toxins like GBCA were still bioactive (Farran et al., 1995). The lack of success of these seed processing methods in improving animal growth or health has strongly indicated the need for genetic approaches to detoxify the common vetch seeds.

### Unsuccessful searches for zero-toxin vetch accessions

Using conventional breeding methods and more recently the use of molecular marker-assisted breeding for genomic selection, plant breeders have prioritized the search for common vetch varieties that have biotic and abiotic stress resistant traits as well as selecting for increasing yield and seed nutritional quality (Francis et al., 2000). However, no concerted effort has been made to select for low or zero GBCA toxin levels in Australian or overseas breeding programs. This has resulted in varieties that are only used by farmers for pasture, green or brown silage or ruminant feed (Francis et al., 2000; Dong et al., 2019; Huang et al., 2019; Mikić et al., 2019). This is mainly due to very limited natural variation in GBCA toxin levels amongst common vetch accessions (Rathjen, 1997). Rathjen et al., screened over 1,700 *V. sativa* accessions and failed to identify a single accession with no GBCA toxin (Rathjen, 1997). Later, screening of a total of 3,000 accessions identified only one line, IR28, with a low (0.3 – 0.4%) GBCA level but not zero-toxin has yet been identified (Ford et al., 2008). Backcrossing IR28 to Jericho white, a spontaneous white flowered mutant of the French commercial variety Languedoc, over 7 generations produced a near homozygous line named Lov 9 (Tate and Searle, unpublished). However, the GBCA levels in the Lov 9 seed from plants grown in shade houses or field conditions ranged from 0.4 – 1.2%, respectively (Tate and Searle, unpublished). These GBCA levels in Lov 9 seed were deemed too high for commercial release of the variety as a low toxin variety. Another strategy to develop a zero GBCA toxin common vetch variety was interspecies crosses of zero toxin species *V. villosa* and *V. pannonica* to common vetch but these resulted in embryo abortion and no viable hybrids were recovered (Searle, unpublished). Considering the limited success to date, other pathways to produce a zero-toxin common vetch variety are required.

### Application of biotechnology to produce zero-toxin vetch

Applications of biotechnology have promised to accelerate crop improvement (Moose and Mumm, 2008). The emergence of new databases, for example genomics, transcriptomics, metabolomics and proteomics information, are now available for most major crops including legumes such as LIS – Legume Information System (Dash et al., 2015) and eFP browser (Patel et al., 2012; Hawkins et al., 2017). By combining the information available in these databases with new bioinformatic tools, we now have the ability to dissect complex traits to determine the underlying gene architecture in a more comprehensive way. In 2018, the 1.8 Gb common vetch genome and transcriptome sequencing projects were initiated at the University of Adelaide, Australia, opening the opportunity to determine the genetic basis of the vetch toxin accumulation. Now we have the tools to identify the genes involved in toxin production and it is possible to examine their functions by overexpressing and mutating candidate genes. Application of CRISPR-Cas (clustered regularly interspaced short palindromic repeats – Cas protein) genome editing to modify agronomically important traits in crops such as wheat, barley, rice and tomato (Liu et al., 2017) will soon be applied to more challenging species including the common vetch. Using CRISPR-Cas genome editing, the nutritional profiles of many crops have been recently demonstrated. For example, in tomato, knocking down genes in the carotenoid metabolic pathway led to a 5-fold increase in lycopene (Li et al., 2018); and in rice, generating mutations in the starch branching enzyme (Berrens et al. 2017) genes increased amylose content by up to 25% and resistant starch to 9.8% (Sun et al., 2017). One of the most significant impacts of CRISPR-Cas genome editing is the potential improvement of a key trait in a commercially released cultivar within 6 months. In contrast conventional breeding of the trait may take 5 – 7 years to release the new cultivar. Importantly, it only takes one generation to obtain an edited plant using genome editing.

A major challenge in non-model plant systems like common vetch, is the delivery of CRISPR-Cas ribonuclear complex into plant cells and subsequent plant regeneration. Unlike crops such as rice and barley where the transformation and plant regeneration systems are standardized (Sahoo et al., 2011; Harwood, 2014), efficient transformation and plant regeneration systems are lacking for common vetch (Ford et al., 2008). Unfortunately, common vetch’s GBCA toxin level have lowered the priority for research and development of these necessary biotechnological tools – for example developing a transformation system. Further investment in common vetch is required to develop transformation and plant regeneration systems to facilitate the application of genome editing for trait improvement.

### Future work and expectations

The environmental benefits, the versatile growth habit and the rich nutritional profile of common vetch makes it an appealing crop to meet future protein food requirements for humans and animals while sustainably contributing to our agricultural system. However, the failure of conventional breeding to develop a zero-toxin common vetch variety requires a new strategy to be employed. The recent availability of new vetch genomic resources and tools for genome editing increase the likelihood of solving the vetch toxicity problem. To make this plausible, a robust and efficient vetch transformation and plant regeneration system are required.

## Author contributions

VN and SR made a substantial contribution and compiled all the authors’ work to prepare for this paper. IRS, SN and IF edited the paper.

## Acknowledgment

The authors would like to thank the Hermon Slade Foundation and The University of Adelaide for the initial funding awarded to IRS and VN, respectively. We thank Jeremy Timmis for useful feedback and editing of the manuscript.

## References

Aguilera, J.F., Bustos, M., and Molina, E. (1992). The degradability of legume seed meals in the rumen: effect of heat treatment. Animal Feed Science and Technology 36, 101–112.

Asgar, M.A., Fazilah, A., Huda, N., Bhat, R., and Karim, A.A. (2010). Nonmeat Protein Alternatives as Meat Extenders and Meat Analogs. Comprehensive Reviews in Food Science and Food Safety 9, 513–529.

Aura, J.E., Carrión, Y., Estrelles, E., and Jorda, G.P. (2005). Plant economy of hunter-gatherer groups at the end of the last Ice Age: plant macroremains from the cave of Santa Maira (Alacant, Spain) ca. 12000–9000 BP. Vegetation History and Archaeobotany 14, 542–550.

Berrens, R.V., Andrews, S., Spensberger, D., Santos, F., Dean, W., Gould, P., Sharif, J., Olova, N., Chandra, T., and Koseki, H. (2017). An endosiRNA-Based Repression Mechanism Counteracts Transposon Activation during Global DNA Demethylation in Embryonic Stem Cells. Cell stem cell 21, 694–703.

Bünemann, E.K., Bongiorno, G., Bai, Z., Creamer, R.E., De Deyn, G., De Goede, R., Fleskens, L., Geissen, V., Kuyper, T.W., and Mäder, P. (2018). Soil quality–A critical review. Soil Biology and Biochemistry 120, 105–125.

Collins, C.L., Henman, D.J., King, R.H., and Dunshea, F.R. (2002). Common vetch (Vicia sativa cv Morava) is an alternative protein source in pig diets. Asia Pacific Journal of Clinical Nutrition 11, S249–S249.

Dash, S., Campbell, J.D., Cannon, E.K.S., Cleary, A.M., Huang, W., Kalberer, S.R., Karingula, V., Rice, A.G., Singh, J., and Umale, P.E. (2015). Legume information system (LegumeInfo. org): a key component of a set of federated data resources for the legume family. Nucleic acids research 44, D1181–D1188.

Dong, R., Shen, S.H., Jahufer, M.Z.Z., Dong, D.K., Luo, D., Zhou, Q., Chai, X.T., Luo, K., Nan, Z.B., Wang, Y.R., and Liu, Z.P. (2019). Effect of genotype and environment on agronomical characters of common vetch (Vicia sativa L.). Genetic Resources and Crop Evolution.

Enneking, D. (1995). The toxicity of Vicia species and their utilization as grain legumes. Centre for Legumes in Mediterranean Agriculture (CLIMA) Occasional Publication No. 6, University ofWestern Australia, NedlandsW. A.

Erskine, W., Smartt, J., and Muehlbauer, F.J. (1994). Mimicry of lentil and the domestication of common vetch and grass pea. Economic Botany 48, 326–332.

Farran, M.T., Uwayjan, M.G., Miski, A.M.A., Sleiman, F.T., Adada, F.A., Ashkarian, V.M., and Thomas, O.P. (1995). Effect of feeding raw and treated common vetch seed (Vicia sativa) on the performance and egg quality parameters of laying hens. Poultry science 74, 1630–1635.

Ford, R., Maddeppungeng, A.M., and Taylor, P.W.J. (2008). Vetch. Compendium of Transgenic Crop Plants.

Francis, C.M., Enneking, D., and El Moneim, A.M.A. (2000). “When and where will vetches have an impact as grain legumes?,” in Linking research and marketing opportunities for pulses in the 21st Century. Springer), 375–384.

Grdc (2018). “Vetch”, in: Grownotes. (Australia: GRDC).

Harper, J.A., and Arscott, G.H. (1962). Toxicity of common and hairy vetch seed for poults and chicks. Poultry Science 41, 1968–1974.

Harwood, W.A. (2014). “A protocol for high-throughput Agrobacterium-mediated barley transformation,” in Cereal Genomics. Springer), 251–260.

Hawkins, C., Caruana, J., Li, J., Zawora, C., Darwish, O., Wu, J., Alkharouf, N., and Liu, Z. (2017). An eFP browser for visualizing strawberry fruit and flower transcriptomes. Horticulture Research 4, 17029.

Henchion, M., Hayes, M., Mullen, A.M., Fenelon, M., and Tiwari, B. (2017). Future Protein Supply and Demand: Strategies and Factors Influencing a Sustainable Equilibrium. Foods 6.

Huang, Y., Li, R., Coulter, J.A., Zhang, Z., and Nan, Z. (2019). Comparative Grain Chemical Composition, Ruminal Degradation In Vivo, and Intestinal Digestibility In Vitro of Vicia Sativa L. Varieties Grown on the Tibetan Plateau. Animals (Basel) 9.

Kaya, A., A. Yörük, M., Esenbuga, N., Temelli, A., and Ekinci, Ö. (2013). Retraction: The Effect of Raw and Processed Common Vetch Seed (Vicia sativa) Added to Diets of Laying Hens on Performance, Egg Quality, Blood Parameters and Liver Histopathology. The Journal of Poultry Science 50, 228–236.

Li, X., Wang, Y., Chen, S., Tian, H., Fu, D., Zhu, B., Luo, Y., and Zhu, H. (2018). Lycopene is enriched in tomato fruit by CRISPR/Cas9-mediated multiplex genome editing. Frontiers in Plant Science 9, 559.

Liu, X., Wu, S., Xu, J., Sui, C., and Wei, J. (2017). Application of CRISPR/Cas9 in plant biology. Acta Pharmaceutica Sinica B.

Lobell, D.B., and Gourdji, S.M. (2012). The influence of climate change on global crop productivity. Plant physiology 160, 1686–1697.

Mao, Z., Fu, H., Nan, Z., and Wan, C. (2015). Fatty acid, amino acid, and mineral composition of four common vetch seeds on Qinghai-Tibetan plateau. Food chemistry 171, 13–18.

Mba, W.P., Longandjo, G.-N.T., Moufouma-Okia, W., Bell, J.-P., James, R., Vondou, D.A., Haensler, A., Fotso-Nguemo, T.C., Guenang, G.M., and Tchotchou, A.L.D. (2018). Consequences of 1.5 C and 2 C global warming levels for temperature and precipitation changes over Central Africa. Environmental Research Letters 13, 055011.

Mikic, A. (2016). Presence of vetches (Vicia spp.) in agricultural and wild floras of ancient Europe. Genetic resources and crop evolution 63, 745–754.

Mikic, A., Mihailović, V., Karagic, Ð., Miloševic, B., Milić, D., Vasiljević, S., Katanski, S., and Živanov, D. (2019). Common vetch (Vicia sativa) multi-podded mutants for enhanced commercial seed production. Proceedings on applied botany, genetics and breeding 180, 78–81.

Moose, S.P., and Mumm, R.H. (2008). Molecular plant breeding as the foundation for 21st century crop improvement. Plant Physiol 147, 969–977.

Navrátilová, A., Neumann, P., and Macas, J. (2003). Karyotype analysis of four Vicia species using in situ hybridization with repetitive sequences. Annals of botany 91, 921–926.

Patel, R.V., Nahal, H.K., Breit, R., and Provart, N.J. (2012). BAR expressolog identification: expression profile similarity ranking of homologous genes in plant species. The Plant Journal 71, 1038–1050.

Pfeffer, M., and Ressler, C. (1967). β-Cyanoalanine, an inhibitor of rat liver cystathionase. Biochemical pharmacology 16, 2299–2308.

Rathjen, J. (1997). The potential of Vicia sativa L. as a grain legume for South Australia. PhD thesis, University of Adelaide.

Reeves, D.W. (1997). The role of soil organic matter in maintaining soil quality in continuous cropping systems. Soil and Tillage Research 43, 131–167.

Ressler, C., Tatake, J.G., Kaizer, E., and Putnam, D.H. (1997). Neurotoxins in a vetch food: Stability to cooking and removal of γ-glutamyl-β-cyanoalanine and β-cyanoalanine and acute toxicity from common vetch (Vicia sativa L.) legumes. Journal of agricultural and food chemistry 45, 189–194.

Roy, D.N., Sabri, M.I., Kayton, R.J., and Spencer, P.S. (1996). β-cyano-L-alanine toxicity: Evidence for the involvement of an excitotoxic mechanism. Natural toxins 4, 247–253.

Sahoo, K.K., Tripathi, A.K., Pareek, A., Sopory, S.K., and Singla-Pareek, S.L. (2011). An improved protocol for efficient transformation and regeneration of diverse indica rice cultivars. Plant methods 7, 49.

Sivakumar, M.V.K., Das, H.P., and Brunini, O. (2005). “Impacts of present and future climate variability and change on agriculture and forestry in the arid and semi-arid tropics,” in increasing climate variability and change. Springer), 31–72.

Sun, Y., Jiao, G., Liu, Z., Zhang, X., Li, J., Guo, X., Du, W., Du, J., Francis, F., and Zhao, Y. (2017). Generation of high-amylose rice through CRISPR/Cas9-mediated targeted mutagenesis of starch branching enzymes. Frontiers in plant science 8, 298.

Tate, M.E., and Enneking, D. (1992). A mess of red pottage. Nature 359, 357–358.

Tate, M.E., Rathjen, J., Delaere, I., and Enneking, D. (1999). Covert trade in toxic vetch continues. Nature 400, 207.

Tenopala, J., González, F.J., and De La Barrera, E. (2012). Physiological responses of the green manure, Vicia sativa, to drought. Botanical Sciences 90, 263–285.

Valentine, S.C., and Bartsch, B.D. (1996). Production and composition of milk by dairy cows fed common vetch or lupin grain as protein supplements to a silage and pasture-based diet in early lactation. Australian journal of experimental agriculture 36, 633–636.

Westhoek, H., Rood, T., Van Den Berg, M., Janse, J., Nijdam, D., Reudink, M., Stehfest, E., Lesschen, J.P., Oenema, O., and Woltjer, G.B. (2011). The protein puzzle: the consumption and production of meat, dairy and fish in the European Union. Netherlands Environmental Assessment Agency.

White, P., Harries, M., Seymour, M., and Burgess, P. (2005). Producing pulses in the northern agricultural region. Western Australia Department of Agriculture and Food.

Who, F.a.O. (2007). “Protein and amino acid requirements in human nutrition. Report of a Joint WHO/FAO/UNU Expert Consultation. WHO Technical Report Series 935”. WHO Geneva (Switzerland)).

